# Patient Subtyping Analysis of Baseline Multi-omic Data Reveals Distinct Pre-immune States Predictive of Vaccination Responses

**DOI:** 10.1101/2024.01.18.576213

**Authors:** Cigdem Sevim Bayrak, Christian Forst, Drew R. Jones, David Gresham, Smruti Pushalkar, Shaohuan Wu, Christine Vogel, Lara Mahal, Elodie Ghedin, Ted Ross, Adolfo García-Sastre, Bin Zhang

**Affiliations:** Department of Genetics and Genomic Sciences, Icahn School of Medicine at Mt Sinai, New York, NY, USA; Mount Sinai Center for Transformative Disease Modeling, Icahn School of Medicine at Mount Sinai, New York, NY, USA; Department of Microbiology, Icahn School of Medicine at Mt Sinai, New York, NY, USA; Department of Biochemistry and Molecular Pharmacology, New York University Langone Health, New York, New York, USA; Center for Genomics and Systems Biology, Department of Biology, New York University, New York, NY, USA; Department of Chemistry, University of Alberta, Edmonton, Alberta, Canada; Systems Genomics Section, Laboratory of Parasitic Diseases, NIAID, NIH, Bethesda, MD, USA; Center for Vaccines and Immunology, University of Georgia, Athens, GA, USA; Department of Infectious Diseases, University of Georgia, Athens, GA, USA; Department of Microbiology, Icahn School of Medicine at Mount Sinai, New York, NY, USA; Global Health and Emerging Pathogens Institute, Icahn School of Medicine at Mount Sinai, New York, NY, USA; Department of Pathology, Molecular and Cell-Based Medicine, Icahn School of Medicine at Mount Sinai, New York, NY, USA; Department of Medicine, Division of Infectious Diseases, Icahn School of Medicine at Mount Sinai, New York, NY, USA; The Tisch Cancer Institute, Icahn School of Medicine at Mount Sinai, New York, NY, USA

**Keywords:** influenza, vaccine, antibody response, molecular subtyping, multi-omic integration

## Abstract

Understanding the molecular mechanisms that underpin diverse vaccination responses is a critical step toward developing efficient vaccines. Molecular subtyping approaches can offer valuable insights into the heterogeneous nature of responses and aid in the design of more effective vaccines. In order to explore the molecular signatures associated with the vaccine response, we analyzed baseline transcriptomics data from paired samples of whole blood, proteomics and glycomics data from serum, and metabolomics data from urine, obtained from influenza vaccine recipients (2019-2020 season) prior to vaccination. After integrating the data using a network-based model, we performed a subtyping analysis. The integration of multiple data modalities from 62 samples resulted in five baseline molecular subtypes with distinct molecular signatures. These baseline subtypes differed in the expression of pre-existing adaptive or innate immunity signatures, which were linked to significant variation across subtypes in baseline immunoglobulin A (IgA) and hemagglutination inhibition (HAI) titer levels. It is worth noting that these significant differences persisted through day 28 post-vaccination, indicating the effect of initial immune state on vaccination response. These findings highlight the significance of interpersonal variation in baseline immune status as a crucial factor in determining vaccine response and efficacy. Ultimately, incorporating molecular profiling could enable personalized vaccine optimization.

## 1 Introduction

Vaccination is a powerful prevention tool against infectious diseases including seasonal influenza (IAV) causing millions of severe illnesses and hundreds of thousands of deaths worldwide each year [1]. Due to antigenic drift, an annual vaccination strategy has been the standard approach for targeting the circulating influenza virus strains. However, it is well-documented that the response to vaccination varies widely among individuals potentially limiting the efficacy of the annual vaccination public health approach [2].

Recent research has revealed several potential biomarkers associated with influenza vaccine response, including pre-existing immunity, age, sex, and obesity [3–6], and further studies examined transcriptomics, glycomics, proteomics, and metabolomics data, which showed their potential links with response to the influenza vaccine [7–10]. The mechanism of vaccine response is complex, and hence not well understood to date. We aimed to extend these efforts on understanding baseline pre-immunity differences, and we hypothesized that the integration of multi-model data may help to characterize baseline molecular subtypes associated with different immune responses to vaccination.

Molecular subtyping is a promising approach for understanding disease progression and responses to interventions [11–14]. It has been used with great success in cancer and neurological diseases such as Alzheimer’s disease and Parkinson’s disease to classify disease subtypes. For this study, it was of interest to investigate molecular subtyping in baseline multi-omic data obtained from a vaccination cohort. To avoid confusion, we hereinafter use the term baseline molecular subtype (BMS) to refer the clusters identified in baseline data of a non-disease cohort. Molecular subtyping of influenza vaccine responses could facilitate development of more effective prevention strategies. To identify baseline biomarkers that can predict vaccine response, we integrated multi-omics data from pre-vaccination biospecimens. We utilized transcriptomics, proteomics, glycomics, and metabolomics data from whole blood, serum, and urine samples, respectively, obtained from a cohort of influenza vaccine recipients. We examined top differential features from each modality contributing to BMS clusters, such as differentially expressed genes, proteins, glycans, and metabolites, and performed pathway enrichment analyses to reveal divergent immune response-related pathways. Ultimately, these findings could offer insights into high-risk individuals susceptible to poor protection, and enable design effective vaccines.

## 2 Materials and methods

### 2.1 Vaccine cohort

Participants were enrolled at the University of Georgia Clinical and Translational Research Unit (Athens, GA, USA) (IRB #3773) from September 2019 to February 2020 (UGA4). All volunteers were enrolled with written, informed consent and excluded if they already received the seasonal influenza vaccine. All participants received a Fluzone® (Sanofi Pasteur) seasonal inactivated influenza vaccine. Most participants received a quadrivalent, standard dose formulation made up of 15 mcg HA per strain of A/H1N1 (A/Brisbane/02/2018), A/H3N2 (A/Kansas/14/2017), B/Yamagata (B/Phuket/3073/2013), and B/Victoria (B/Colorado/6/2017-like strain). Some of the older participants (≥65) chose the high-dose vaccine, which was a trivalent composition lacking a B/Yamagata strain but formulated with 60 mcg HA/strain of the others.

### 2.2 Multi-omic data

In this study, we used RNAseq data from whole blood samples of 275 subjects (12,583 genes), proteomics (273 proteins) and glycomics data (probes including 68 lectins and 14 carbohydrate-binding antibodies) from serum samples of 151 subjects, and urine metabolomics data (15,903 metabolites) of 179 subjects from the UGA4 cohort. The number of subjects with all four data types available was 62 (Figures 1A and 1B). Combined multi-omic data is available at https://www.synapse.org/#!Synapse:syn52749029.

**Figure 1.**
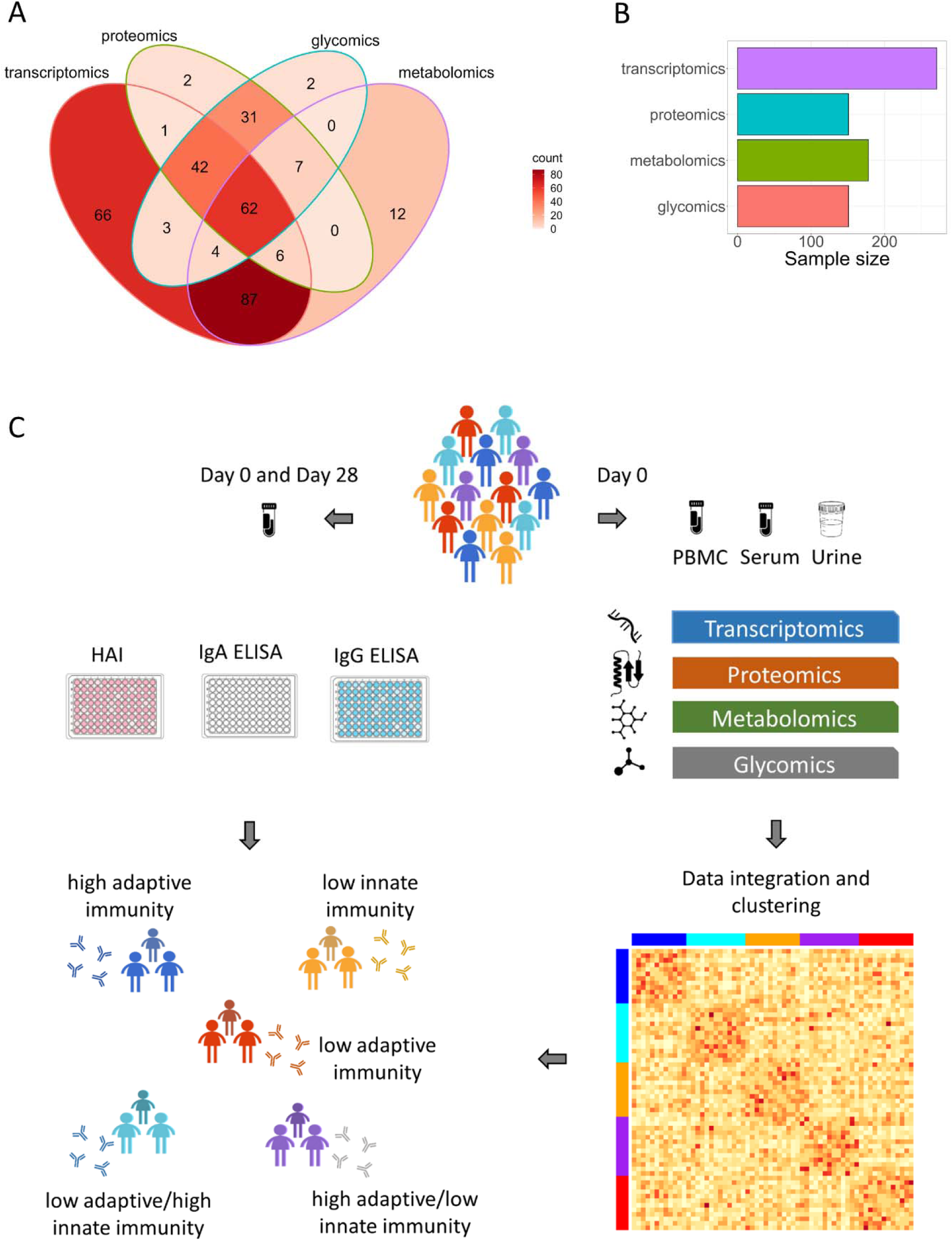
Dataset and main steps of the analytical procedure used in the study. (A) Venn diagram showing the number of samples with single or multi-omics data in the UGA4 cohort. Number of samples with all four omic data type is 62. (B) Number of samples in the UGA4 cohort by data modality. (C) Procedure to identify baseline molecular subtypes from pre-vaccination multi-omics data.

#### 2.2.1 Transcriptomics

RNA sequencing data obtained from 275 whole blood samples from the UGA4 study was published [7]. Each tube contained 2.5 mL of blood that was collected prior to vaccination (day 0). Sequence reads were aligned to the human transcriptome using the nf-core/rnaseq nextflow pipeline [15] with default parameters. This pipeline aligned reads to the human genome (GRCh37) using STAR [16] followed by BAM-level quantification with Salmon [17] to generate a feature counts table. Read counts within samples, and between samples were then normalized and filtered using the edgeR (v3.40.2) R-package [18]. Four outlier samples were filtered out based on plotMDS function from the limma (v3.54.2) R package [19]. Logarithm of TPM (transcript per million) values were calculated and adjusted for batch, age, sex, race, BMI, month of vaccination, and prior vaccination status by fitting a linear model. The data further went through Z transform to normalize the profile of each gene and the expression values were restricted to the range of minus and plus 4 standard deviations.

#### 2.2.2 Proteomics

The details of 225 serum proteome probes preparation from 160 influenza vaccine recipients was previously published [9]. All 225 raw files were first converted to the HTRMS format with the HTRMS converter. The converted files were then analyzed with the directDIA method using default settings. Data preprocessing includes quality control, batch correction, preprocessing by mapDIA, and normalization as described previously [9, 20]. We further processed the data by adjusting for age, sex, BMI, month of vaccination, and prior vaccination status, and then applied Z-transform.

#### 2.2.3 Glycomics

The glycosylation data including 68 lectins and 14 carbohydrate-binding antibodies from 160 subjects was previously published [8]. Glycomic analysis was performed using a dual-color lectin microarray technology as previously described. The details of fluorescent labeling of serum proteins and lectin microarray generation can be found in [8]. We then adjusted the data for age, sex, BMI, month of vaccination, and prior vaccination status, and then applied Z-score transform.

#### 2.2.4 Metabolomics

The data from the analysis of urine samples from 179 subjects including 15,903 putatively identified metabolites was previously published [10]. The details of metabolite extraction, batch design, and quality control were previously described. Before the downstream analysis, data was further adjusted for age, sex, race, BMI, month of vaccination, and prior vaccination, and normalized by Z-score transform.

### 2.3 HAI assay

Blood samples were collected from participants prior to vaccination (day 0) and again post vaccination on day 28. Hemagglutinin inhibition (HAI) assays against each vaccine strain were performed with serum from each participant on day 0 and day 28.

### 2.4 Definition of Antibody Responses

The vaccine response was assessed by hemagglutination inhibition assay (HAI) and enzyme-linked immunosorbent assay (ELISA) immunoglobulin G (IgG) and immunoglobulin A (IgA) antibody levels. For each vaccine strain, we calculated the response scores by calculating the logarithm of fold changes between day 0 and day 28 antibody titer levels. Then, we calculated average response values. Specifically, the composite baseline and composite day 28 values were calculated by taking the summation of 3 (high dose) or 4 (standard dose) strains depending on the administered vaccine dose. Following the previous seroconversion definition [3], we have then calculated the seroconversion as the logarithm of fold change in total antibody levels on day 28 compared to the baseline values (seroconversion = log_2_(*D*28/*D*0)). All response values were then adjusted for age, sex, race, BMI, month of vaccination, and prior vaccination.

### 2.5 Data integration and subtype identification

To integrate multi-omic data modalities, we have used the similarity network fusion (SNF) model [21]. For a given cohort, the SNF model first computes patient similarity networks obtained from each data type, and then fuses them into one network representing the full spectrum of underlying data. We have used the SNFtool R package (v2.3.1) with parameter settings number of neighbors K=20, total number of algorithmic iterations T=20, and the hyper-parameter alpha=0.5. The distance matrices were calculated using Euclidean distance for each data type. The affinity matrices were then generated representing the neighborhood graph based on the given parameters K and alpha. The SNF function was used to integrate the affinity matrices. The number of clusters were estimated (2 to 5 clusters) by the eigen-gap and rotation cost methods. To predict the baseline molecular subtypes (BMS), we used the default unsupervised clustering method of the SNFtool, spectral clustering, on the fused matrix.

### 2.6 Differential Gene Expression Analysis

To identify differentially expressed genes (DEGs) of each BMS, we used the EdgeR (v3.40.2) in R. Low-expressed genes were filtered out using the filterByExpr() function. Then, the remaining counts were normalized, and dispersions were estimated and fit by glmQLFit() function which is a quasi-likelihood negative binomial generalized log-linear model.

### 2.7 Differentially Expressed Proteins, Metabolites, and Glycomes

For each molecule, we have calculated average expression levels of each BMS group. Then we have determined the logarithm of fold change by dividing the average expression of one BMS to the average expression level of other BMSs, and performed Kruskal-Wallis test to identify statistical significance of each expression.

### 2.8 Pathway Analysis

We used GOtest (v1.0.9) R package to perform pathway enrichment analysis using the MSigDB collections [22]. The query genes were the DEGs determined by comparing each BMS to average of other four BMSs and filtered by a Pvalue cutoff of 0.05 and fold change cutoff of 1.5.

For the pathway analysis of differentially expressed proteins (DEPs), we have used the ReactomePA (v1.42.0) R package where the DEPs were selected with a Pvalue cutoff of 0.05 and fold change cutoff of 1.2.

We have selected the top differentially expressed metabolites with a Pvalue cutoff of 0.05 and fold change cutoff of 2 and used MetaboAnalyst web-based platform to perform enrichment analysis [23].

### 2.9 Statistical Analysis

To analyze associations between subtypes and clinical features as well as day 0 and day 28 antibody levels, we performed Kruskal-Wallis test. To compare features of each BMS pair, we used ggsignif (v0.6.4) R package to calculate Wilcoxon test and to add annotations to the violin plots.

The significance of overlapping DEG sets were calculated by the SuperExact (v1.1.0) R package [24].

The top subtype-specific features were determined using Kruskal-Wallis one-way ANOVA test by comparing Z-transformed expression levels of each BMS to the other four BMSs (i.e., one vs others).

## 3 Results

### 3.1 Integrated four-modal data identifies five baseline molecular subtypes

To determine potential baseline molecular subtypes (BMS), we applied clustering on integrated baseline transcriptomics, proteomics, glycomics, and metabolomics data from 62 participants using the SNF method. Figure 1C illustrates the clustering analysis pipeline. The optimal number of clusters was estimated as k=5 according to rotation cost (for k=2:5) and eigengap heuristic tests. The five clusters, baseline molecular subtypes, were named by their molecular properties: (i) high adaptive immunity subtype (S-HiAI), (ii) low adaptive/high innate immunity subtype (S-LoAI/HiII), (iii) low innate immunity subtype (S-LoII), (iv) high adaptive/low innate immunity subtype (S-HiAI/LoII), and (v) low adaptive immunity subtype (S-LoAI). To explore associations between different subtypes and the vaccination response, we performed the Kruskal-Wallis test. The predicted BMSs were significantly associated with the logarithm of A/H3N2/KA17 IgA levels on day 0 and day 28, and logarithm of B/Yamagata/Phu13 HAI level on day 28 (BH-adjusted P=0.02).

Demographic characteristics of 62 study participants grouped by BMS are given in Table 1. The subtypes were equally distributed across all demographic and clinical features, demonstrating that the clusters do not represent physiological differences.

**Table 1.**
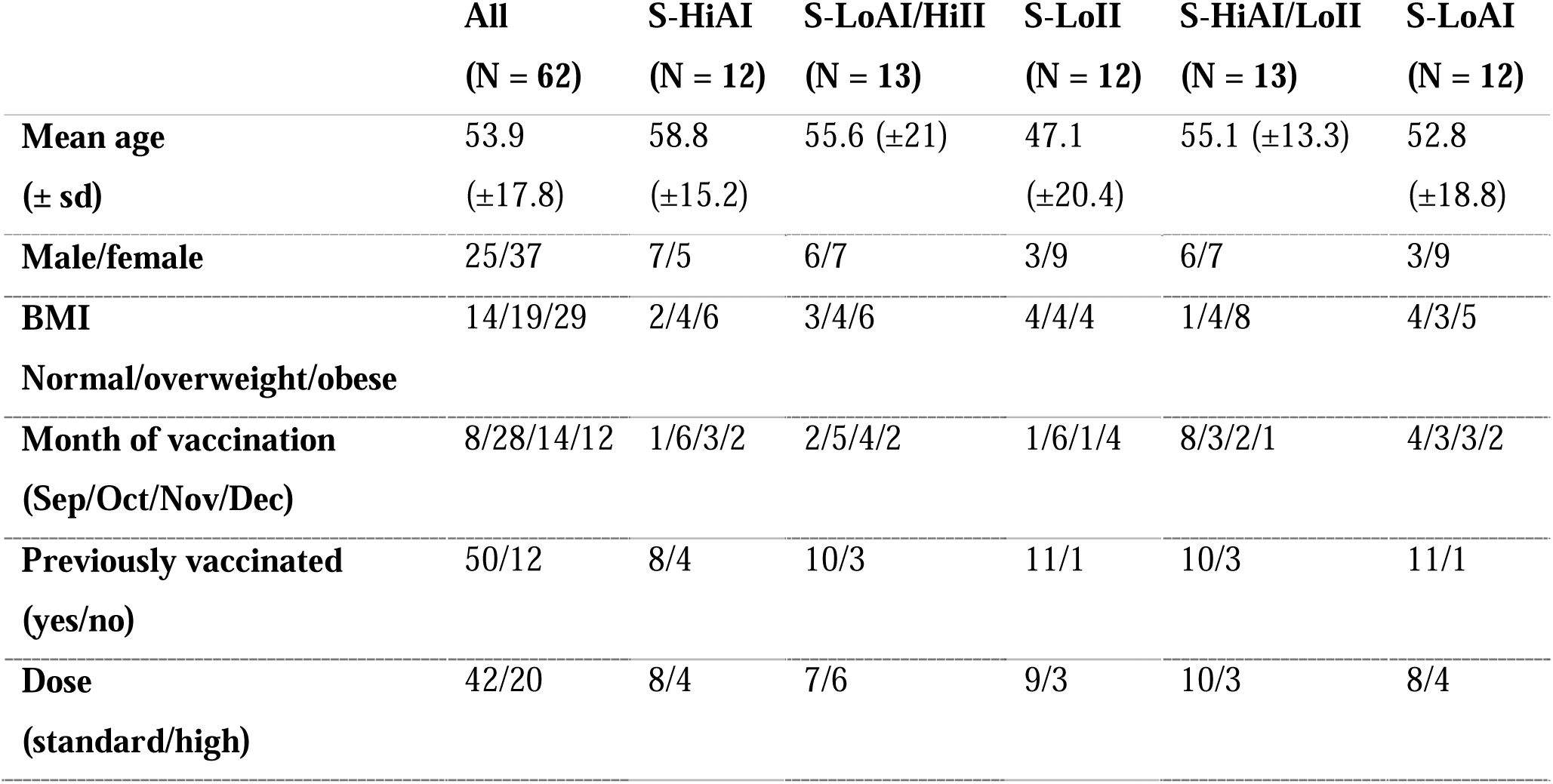
Characteristics of study participants stratified by predicted baseline molecular subtypes.

### 3.2 Subtype-specific molecular functions and vaccination response

To characterize subtype-specific molecular features, we applied pathway enrichment analysis of differentially expressed genes for each BMS compared to the other BMSs. Each BMS showed distinct molecular signatures with subtype-specific up- and down-regulated pathways, suggesting heterogeneous responses to vaccination. Figure 2 illustrates top enriched GO terms for each BMS using the Molecular Signatures Database (MSigDB) collection [22]. We further evaluated differential metabolite levels of the predicted subtypes by performing metabolite pathway enrichment analysis. We found that the differentially expressed metabolites of each BMS were significantly enriched in different KEGG pathways. Similarly, we identified the pathways of differentially expressed proteins. Due to lack of a straightforward computational approach for identifying enriched pathways from serum glycomes, we did not investigate glycoprotein pathways.

**Figure 2.**
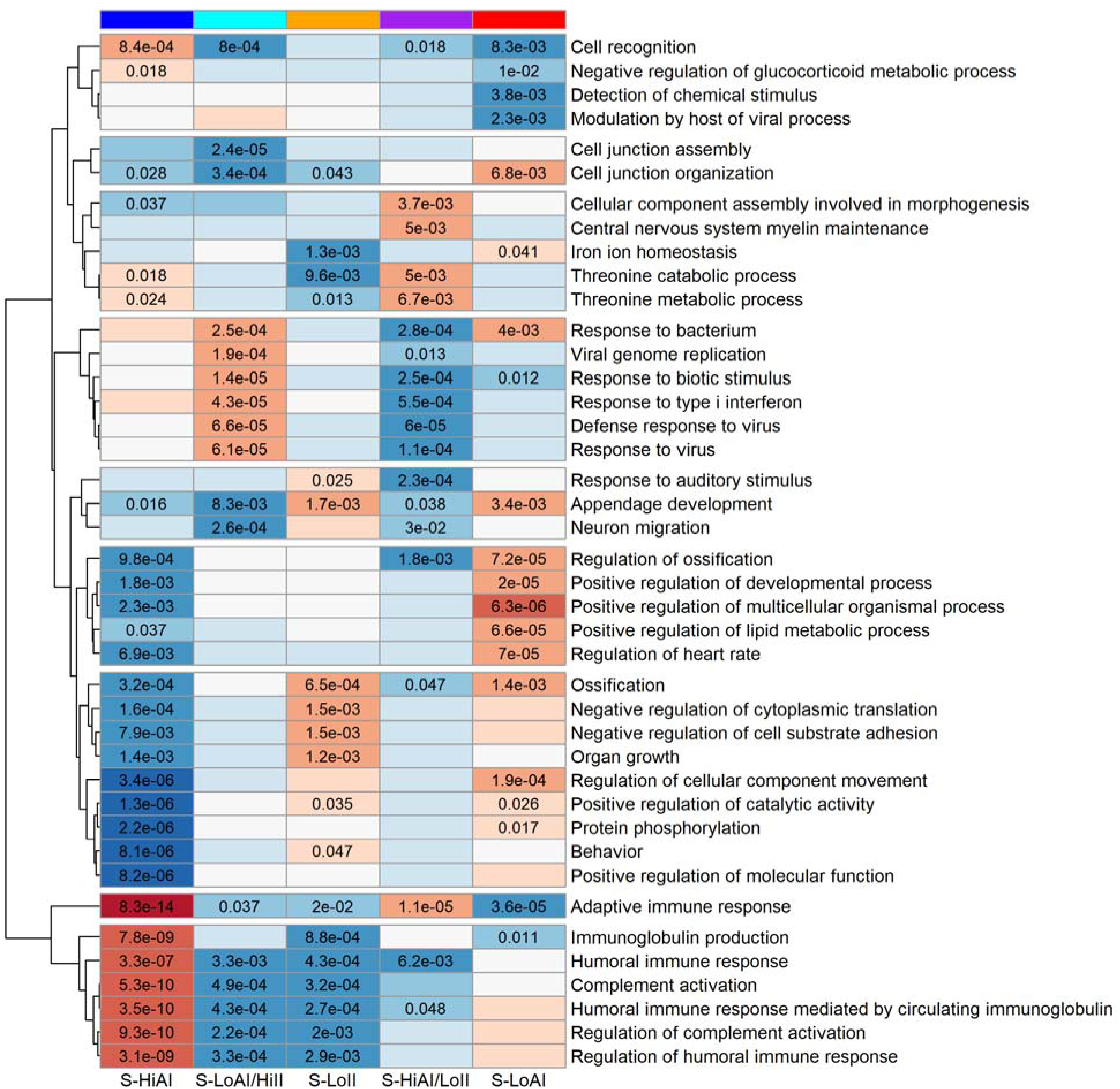
Pathway enrichment analysis of differentially expressed molecules of each BMS. MSigDB GO-BP enrichment analysis of differentially expressed genes identified in each subtype. Columns represent the baseline molecular subtypes: S-HiAI (high adaptive immunity subtype), S-LoAI/HiII (low adaptive/high innate immunity subtype), S-LoII (low innate immunity subtype), S-HiAI/LoII (high adaptive/low innate immunity subtype), and S-LoAI (low adaptive immunity subtype). Hierarchical clustering was applied on the rows to illustrate pathway clusters. Red color indicates up-regulated pathways, blue color represents down-regulated pathways, and the shades indicate the significance.

To better understand differences in vaccination responses, we also examined the associations between BMS groups and antibody levels pre- and post-vaccination. It is noteworthy that although some of the subtypes and pre- and/or post-vaccination antibody levels showed significant associations, none of the subtypes were found associated with vaccination response. In this study, all antibody values were adjusted for age, sex, race, BMI, month of vaccination, and prior vaccination.

Below, we present the molecular features and vaccination response of each predicted baseline molecular subtype in detail.

*S-HiAI (high adaptive immunity subtype):* We observe that the DEGs involved in adaptive and humoral immune responses, immunoglobulin production and B cell mediated immunity were significantly up-regulated in the baseline blood transcriptomic data of the S-HiAI subtype (Figures 3A-3C). These up-regulated DEGs include immunoglobulin heavy (IGH), the immunoglobulin kappa (IGK), and the immunoglobulin lambda (IGL) genes, as well as T-cell receptor alpha (TRA), T-cell receptor beta (TRB) genes. Humoral immunity (antibody-mediated immunity) is a type of adaptive immune response, which is specific to the pathogen. In the antibody response, B cells are activated to produce antibodies that bind to the antigens and neutralize them. Results demonstrate that the S-HiAI subtype has higher pre-existing adaptive immunity than the other subtypes.

**Figure 3.**
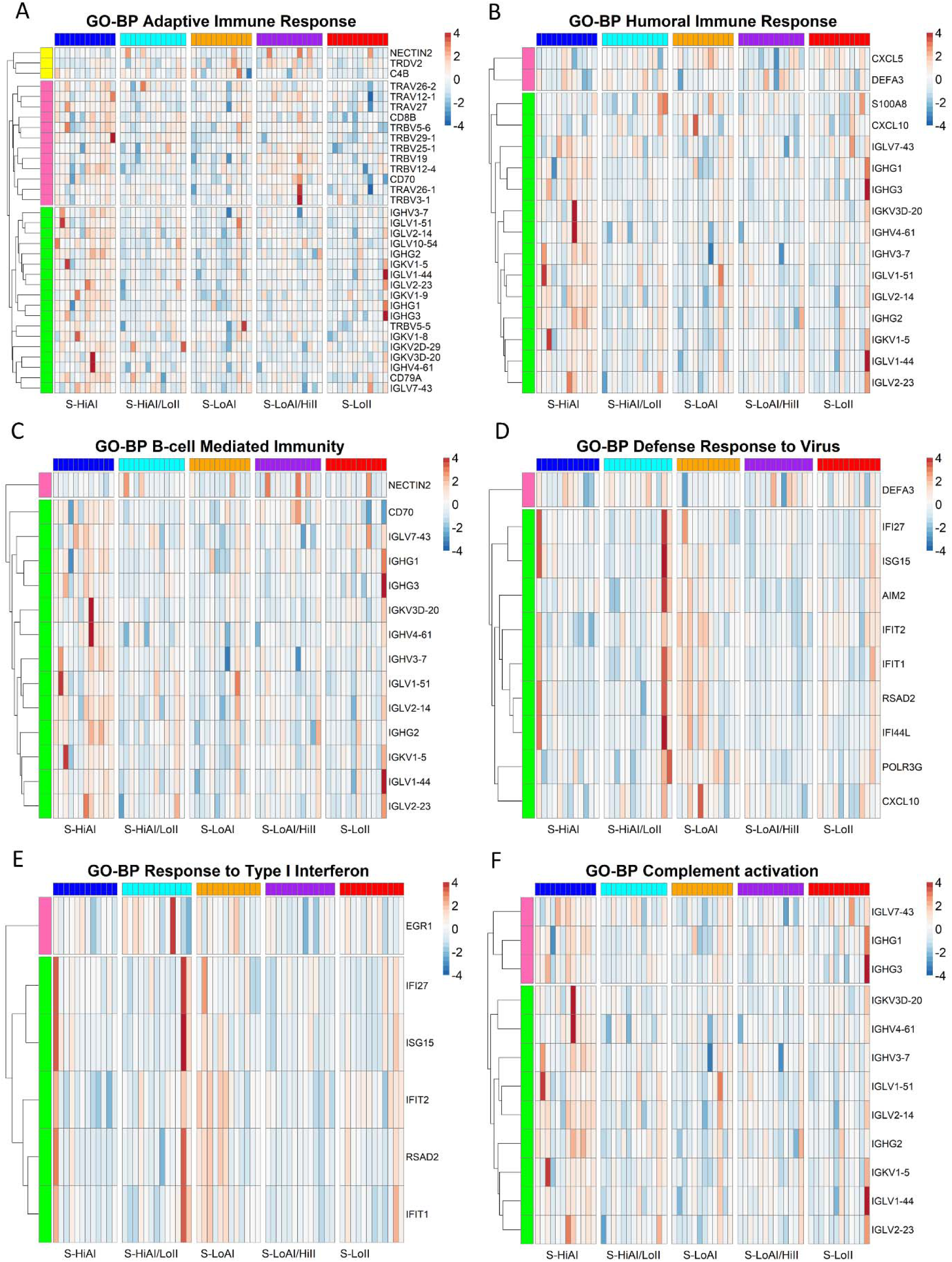
Biological processes highlighting distinct molecular features of baseline molecular subtypes. The colored horizontal bars on top represents the baseline molecular subtypes: S-HiAI (high adaptive immunity subtype), S-LoAI/HiII (low adaptive/high innate immunity subtype), S-LoII (low innate immunity subtype), S-HiAI/LoII (high adaptive/low innate immunity subtype), and S-LoAI (low adaptive immunity subtype). The rows correspond to DEGs identified in each subtype. The expression values are Z-transformed logCPM values. Hierarchical clustering was applied on the rows to illustrate gene clusters. (A) DEGs of GO-BP adaptive immune response pathway, (B) DEGs of GO-BP humoral immune response, (C) DEGs of GO-BP B cell mediated immunity, (D) DEGs of GO-BP defense response to virus, (E) DEGs of GO-BP response to type I Interferon, (F) DEGs of GO-BP complement activation.

Based on the metabolite pathway enrichment analysis, arginine biosynthesis was significantly enriched in the S-HiAI subtype. Up-regulated arginine biosynthesis indicates active immune response and inflammation [25–27]. In the DEPs of the S-HiAI subtype, platelet degranulation, and synthesis of 5-eicosatetraenoic acids were among up-regulated pathways. Platelets are known as key players in inflammation [28]. 5-eicosatetraenoic acid is a metabolite of arachidonic acid produced by 5-lipoxygenase which are central players of inflammatory pathways [29]. These findings support high existing immunity.

The average IgA level of the S-HiAI subtype was significantly lower than of the S-LoAI/HiII, S-HiAI/LoII, and S-LoAI subtypes both pre-vaccination and post-vaccination (Figure 4A). More specifically, the A/H3N2 IgA level was significantly lower in the S-HiAI subtype at day 0, which remained low after vaccination (Figure 4B). The only significant antibody level change post-vaccination was for B/Yamagata HAI with an increase in the S-HiAI subtype (Figure 4C). This subtype is characterized by low IgA antibody levels in the blood and up-regulated humoral immunity pathways indicating increased activity of B cells and antibody production at the gene level. While this is representative of complex immune dynamics, it suggests that either the humoral immune process has not yet yielded higher circulating IgA or that blood IgA levels do not yet correspond to antibody stimulation occurring in mucosal tissues, such as those found in the respiratory tract.

**Figure 4.**
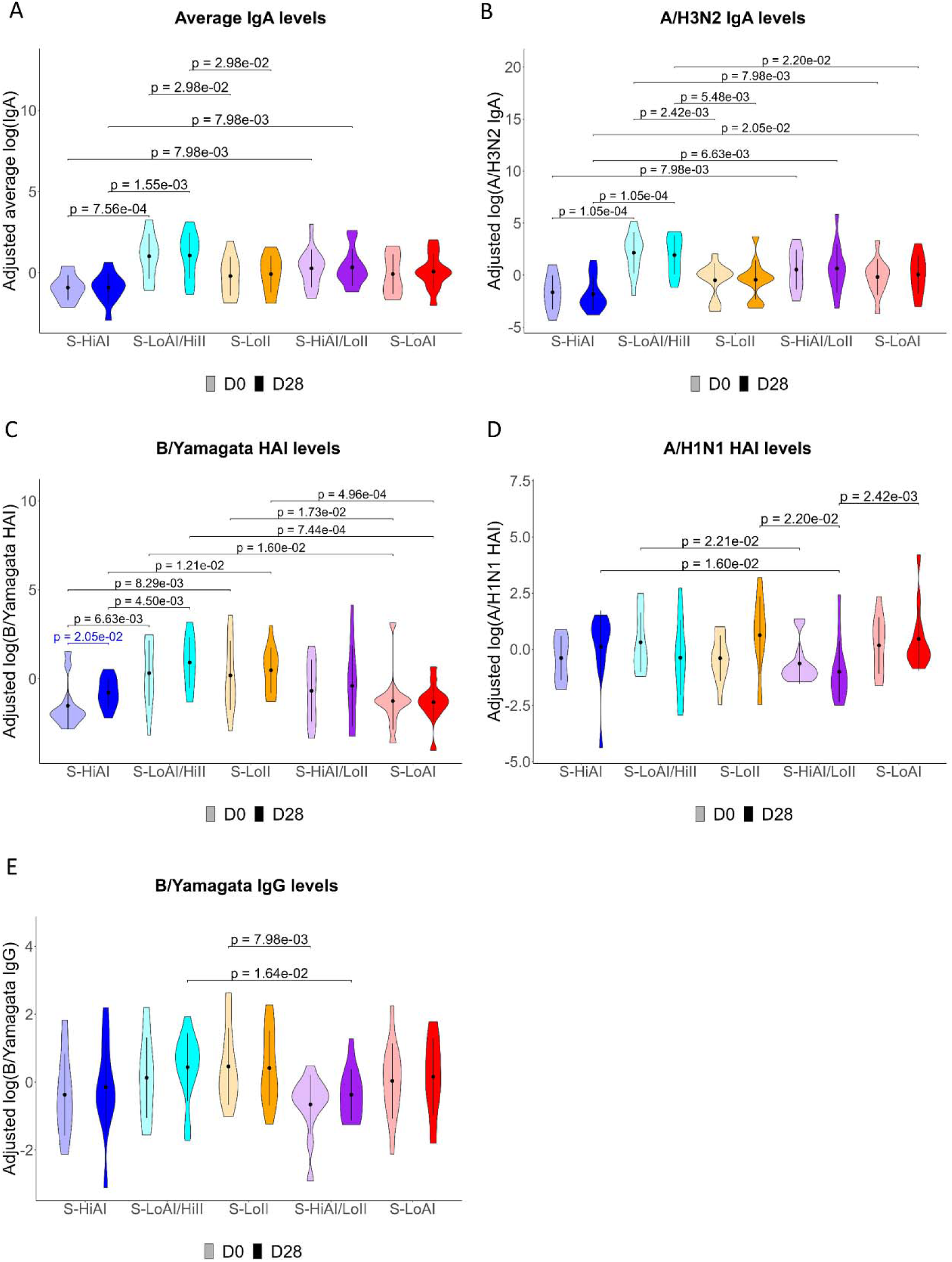
Pre- and post-vaccination antibody levels across baseline molecular subtypes. The x-axis shows the predicted baseline molecular subtypes: S-HiAI (high adaptive immunity subtype), S-LoAI/HiII (low adaptive/high innate immunity subtype), S-LoII (low innate immunity subtype), S-HiAI/LoII (high adaptive/low innate immunity subtype), and S-LoAI (low adaptive immunity subtype), and the y-axis shows the covariate adjusted antibody levels measured on day 0 and day 28. The pairwise comparisons were performed by the Wilcoxon test.

*S-LoAI/HiII (low adaptive/high innate immunity subtype):* Defense response to virus, response to type I interferon, innate immune response, and interferon alpha/beta signaling pathways were up-regulated in the S-LoAI/HiII subtype (Figures 3D and 3E), while humoral immune response and complement activation were down-regulated. The up-regulated DEGs of the S-LoAI/HiII subtype includes interferon-inducible genes (e.g., *IFI27*, *IFI44*, *IFI44L*, and *IFIT1*). Interferons provide natural defense and are released by host cells in response to viral infections. Together, these findings indicate that the S-LoAI/HiII subtype is characterized by high innate but low adaptive pre-existing immunity. This may indicate a pre-existing infection modulating innate immune mediators to actively suppress adaptive immunity [30].

The metabolite pathways alanine-aspartate-glutamate metabolism, arginine biosynthesis, glycine-serine-threonine metabolism, and tryptophan metabolism were enriched in the metabolomics data of the S-LoAI/HiII subtype. With respect to proteins, ficolins serve as recognition subunits of the innate immune system, through the binding of carbohydrate arrays on the surfaces of pathogens and trigger complement activation via the lectin pathway [31, 32]. The pathways related to activation of complement by lectin and binding of ficolin were among the significantly down-regulated pathways of DEPs of the S-LoAI/HiII subtype.

The A/H3N2 IgA level of the S-LoAI/HiII subtype was significantly higher than the other subtypes both pre- and post-vaccination (Figure 4B). Moreover, B/Yamagata HAI levels of the S-LoAI/HiII subtype were higher than in the S-HiAI and S-LoAI subtypes on both day 0 and day 28 (Figure 4C). High circulating IgA antibodies specific to H3N2 influenza coupled with molecular evidence of enhanced antiviral/inflammatory pathways suggest the immune system is actively responding to an H3N2 infection or exposure in this subtype. This is consistent with an active (or very recent) influenza infection rather than a baseline immunological state.

*S-LoII (low innate immunity subtype):* Immunoglobulin production, humoral immune response, and complement activation were among the significantly down-regulated pathways of the S-LoII subtype (Figures 3B and 3F). The canonical pathways related to activation and regulation of the immune response such as complement cascade, initial triggering of complement, CD22 mediated BCR regulation, creation of C4 and C2 activators were also down-regulated, suggesting low innate immunity. Down-regulated DEGs involved in these pathways included immunoglobulin genes (e.g., *IGHV4-61*, *IGKV3D-20*, *IGLV1-44*, and *IGLV7-43*). Immunoglobulins play an important role in the immune system by recognizing and defending against pathogens. The genes related to immunoglobulin production were expressed at significantly lower levels in the S-LoII subtype compared to the others. This indicates some kind of immune dysfunction or impaired antibody-mediated immune response.

With respect to metabolites, glycine-serine-threonine metabolism, caffeine metabolism, which is known to be associated with immune response [33, 34], and vitamin B6 metabolism, which has been implicated in the regulation of the immune response [35, 36], were enriched in the S-LoII subtype. Based on the DEPs, complement cascade-related pathways were down-regulated, supporting low innate immunity.

*S-HiAI/LoII (high adaptive/low innate immunity subtype):* As opposed to the S-LoAI/HiII subtype (low adaptive/high innate immunity subtype), pathways including defense response to virus, response to type I interferon, innate immune response, and interferon alpha/beta signaling were down-regulated in the S-HiAI/LoII subtype (Figures 3D and 3E). Interestingly, adaptive immune response was up-regulated while humoral immune response was down-regulated (Figure 3A and 3B). Two main classes of the adaptive immune response are antibody-mediated (humoral) and cell-mediated responses which are carried out by B cells and T cells, respectively. Up-regulated DEGs of the S-HiAI/LoII subtype included T-cell receptor genes (e.g., *TRAV26-1*, *TRBV19*, *TRBV25-1*, *TRBV3-1*, and *TRDV2*) and down-regulated DEGs included interferon-inducible genes (e.g., *IFI27*, *IFI44L*, *IFIT1*, and *IFIT2*) and immunoglobulin genes (e.g., *IGHG3*, and *IGHV4-61*). Together, these results indicate imbalance between cellular, cytokine, and humoral responses suggesting that the S-HiAI/LoII subtype has overactive cellular (T cell mediated adaptive immunity) but underactive innate immune responses.

Alanine-aspartate-glutamate metabolism, arginine biosynthesis, and glycine-serine-threonine metabolism were enriched metabolite pathways of the S-HiAI/LoII subtype. Complement cascade, which is a part of the immune system, was significantly up-regulated in the DEPs of the S-HiAI/LoII subtype [37]. Lipoprotein related pathways including chylomicron assembly and remodeling, and plasma lipoprotein assembly were significantly down-regulated. Lipoproteins have been suggested to have a role in innate immunity as they may prevent infections [38].

A/H1N1 HAI levels of the subtypes have changed in different directions after vaccination when comparing different clusters against each other (Figure 4D). Although, the changes were not statistically significant, A/H1N1 HAI levels of S-LoAI/HiII and S-HiAI/LoII subtypes were lower on day 28 compared to day 0, and it were higher after vaccination in other subtypes. As a result, we observe that the S-HiAI/LoII subtype had significantly lower A/H1N1 HAI levels compared to the other subtypes (i.e., S-HiAI, S-LoII, and S-LoAI) post-vaccination. Lower H1N1 HAI titers may reflect reduced neutralizing antibody levels against the flu strain. While cellular immune activation is high, the antibody-mediated humoral response seems to be defective in the S-HiAI/LoII subtype.

*S-LoAI (low adaptive immunity subtype):* Contrary to the S-HiAI subtype (high adaptive immunity subtype), adaptive immune system and immunoglobulin production were down-regulated pathways in the S-LoAI subtype compared to the other subtypes (Figure 3A). Up-regulated pathways of the S-LoAI subtype on the other hand, included regulation of biological processes such as multicellular organismal process, developmental process, lipid metabolic process, and regulation of heart rate. Lipid metabolism plays a major role in the regulation of the immune system [39], and interactions between inflammation and autonomic functions have been discussed in the literature [40]. Up-regulated DEGs of the S-LoAI subtype including integrin subunit genes (ITGs), interleukins (IL), and P2Y receptors, were enriched in the pathways related to regulation of immune responses, such as IL-4, integrin cell signaling, cytokine-cytokine receptor interaction, and P2Y receptors pathways. [41, 42] (Figure 5). Overall, these findings indicate gene expression shift towards innate immune response genes rather than adaptive immune response genes for antigen-specific responses in the S-LoAI subtype.

**Figure 5.**
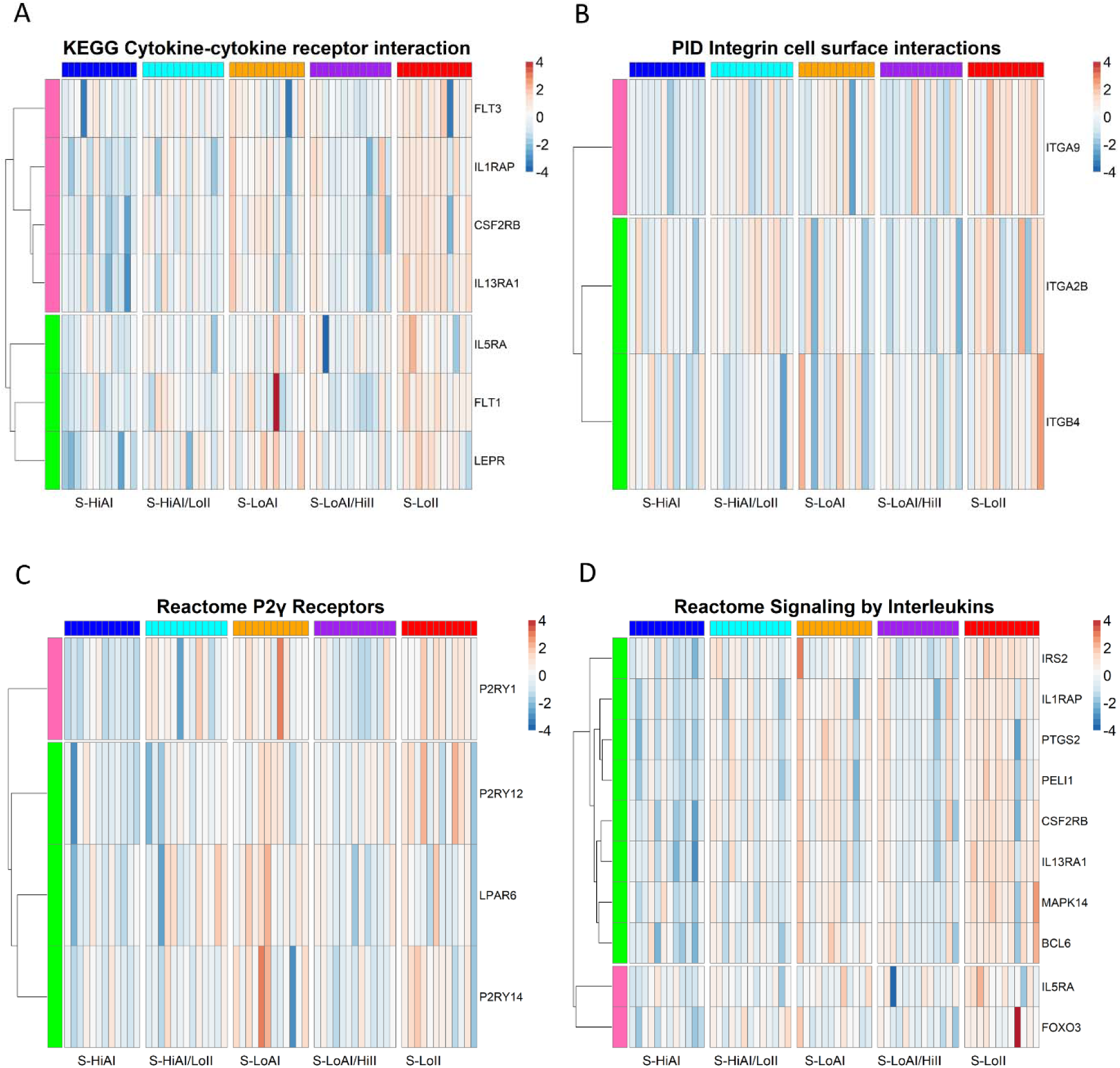
Canonical pathways highlighting distinct molecular features of baseline molecular subtypes. The colored horizontal bars on top represents the baseline molecular subtypes: S-HiAI (high adaptive immunity subtype), S-LoAI/HiII (low adaptive/high innate immunity subtype), S-LoII (low innate immunity subtype), S-HiAI/LoII (high adaptive/low innate immunity subtype), and S-LoAI (low adaptive immunity subtype). The rows correspond to DEGs identified in each subtype. The expression values are Z-transformed logCPM values. Hierarchical clustering was applied on the rows to illustrate gene clusters. (A) DEGs of KEGG Cytokine-cytokine receptor interaction pathway, (B) DEGs of PID Integrin cell surface interactions pathway, (C) DEGs of Reactome P2γ receptors pathway, (D) Reactome Signaling by Interleukins pathway.

Based on the metabolomics data, the pathways including alanine-aspartate-glutamate metabolism, arginine biosynthesis, and glycine-serine-threonine metabolism were all significantly enriched in the S-LoAI subtype. In the DEPs, platelet degranulation and antimicrobial peptides, which are known to participate in innate immunity, were down-regulated [43–45].

Furthermore, we observed that the B/Yamagata HAI level of the S-LoAI subtype was significantly lower than in the S-LoAI/HiII and S-LoII subtypes on day 0, and this difference was even more significant after vaccination on day 28 (Pvalues on D0 = 1.60E-02 and 1.73E-02, Pvalues on D28 = 7.44E-04 and 4.96E-04, respectively) (Figure 4C). Together, lower pre-existing and vaccine-induced Yamagata HAI levels may imply higher susceptibility to infection with this flu strain due to absence of sufficient neutralizing antibody levels.

### 3.3 Gene signatures of the predicted subtypes

To understand genetic features of the subtypes, we compared the top DEGs identified by comparing each subtype with the remaining four subtypes. As shown above, while the DEGs were mostly associated with immune response-related pathways in all subtypes, we observe that some genes (e.g., immunoglobulin genes, interferons, T-cell receptor genes, etc.) were up-regulated in one subtype, while they were down-regulated in the other. For example, DEGs from immunoglobulin family (e.g., *IGHV4-61*, *IGKV3D-20*), which are antigen receptors of the B cells of the adaptive immune response, were up-regulated in the S-HiAI (high adaptive immunity) subtype, and down-regulated in the S-LoAI/HiII, S-LoII, and S-HiAI/LoII subtypes (Figures 6 and 7). T-cell receptor DEGs were up-regulated in the S-HiAI and S-HiAI/LoII subtypes, subtypes with high adaptive immunity, and down-regulated in the S-LoAI (low adaptive immunity) subtype. Further, the genes from interferon family (e.g., *IFI27*, *IFI44L*, *IFITI*), known to be involved in defense response, were up-regulated in the S-LoAI/HiII (low adaptive/high innate immunity) subtype and down-regulated in the S-HiAI/LoII (high adaptive/low innate immunity) subtype. P2Y receptors involved in inflammation response, were up-regulated in the S-LoAI (low adaptive immunity) and S-LoII (low innate immunity) subtypes.

**Figure 6.**
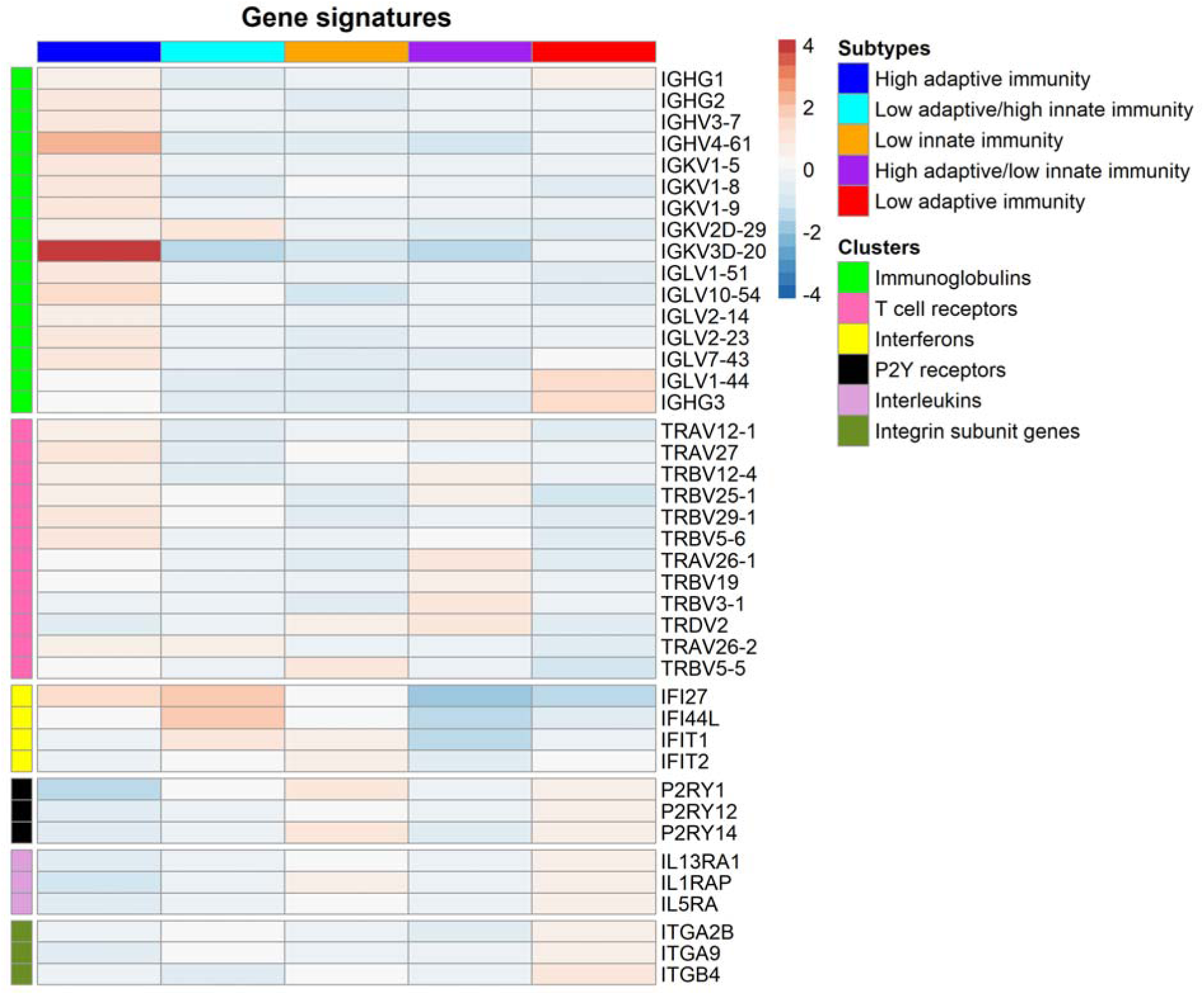
Heatmap of distinct DEG signatures of baseline molecular subtypes. The columns represent the baseline molecular subtypes (BMSs) as indicated by colored bars on top, and the rows indicate top DEGs from immunoglobulin, T cell receptor, interferon, and P2Y receptors. Cell values indicate the logarithm of fold change, which is calculated by comparing one BMS to the average of other four BMSs.

**Figure 7.**
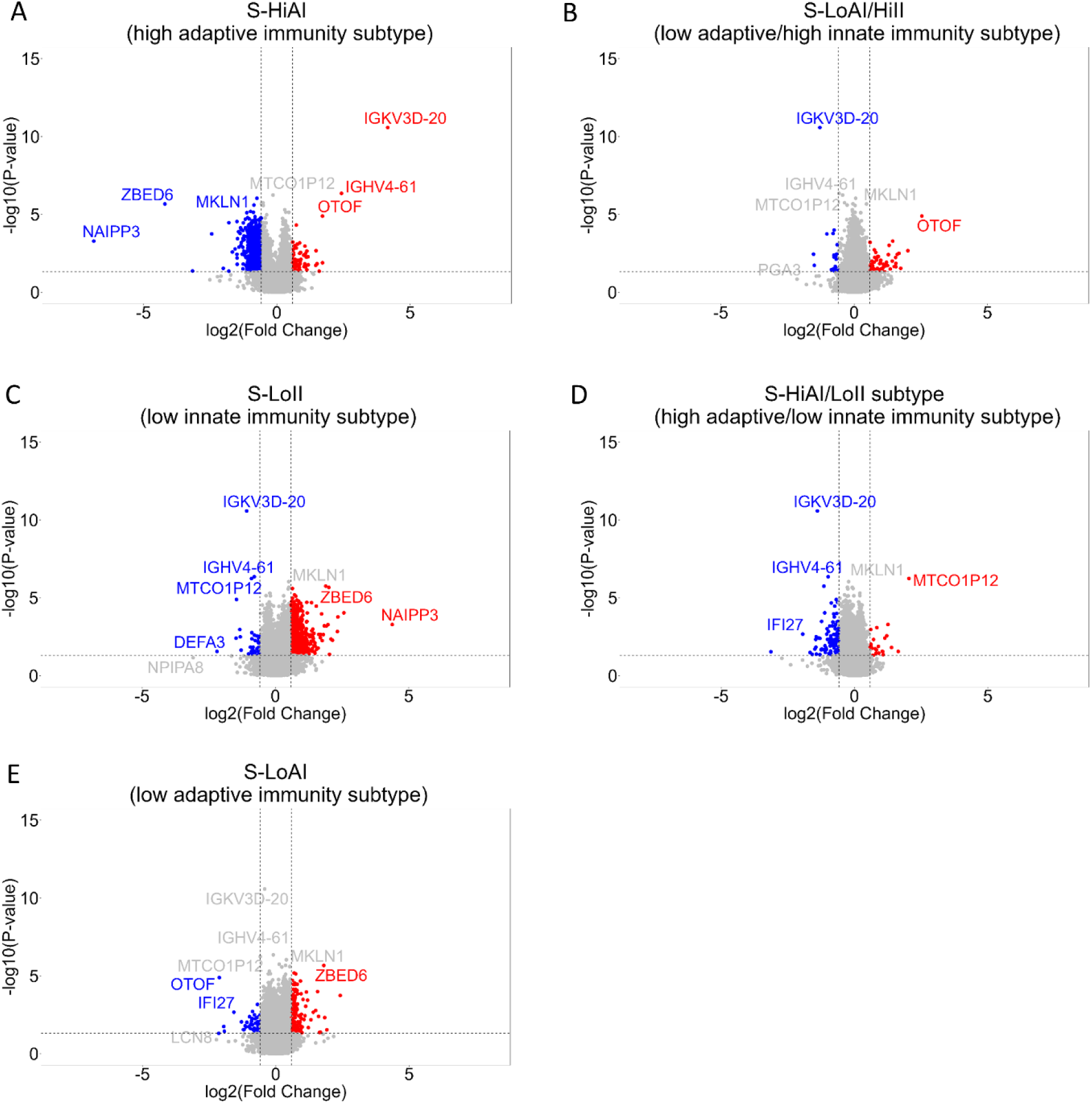
Volcano plots of gene expression values of each BMS compared to the average of other four BMSs. The volcano plots show the fold-change (x-axis) versus the significance (y-axis) of the identified DEGs of each subtype. The vertical and horizontal dotted lines show the cut-off of fold change⍰=⍰⍰±⍰1.2, and of P-value⍰=⍰0.05, respectively.

As seen in Figure 7, differentially expressed genes *OTOF*, and *ZBED6* had opposite directions in different subtypes. Otoferlin gene, *OTOF*, is a type I interferon-induced gene and is a Ca2+ sensor playing a significant role in the synaptic transmission by tethering synaptic vesicles to the plasma membrane [46, 47], This gene was up-regulated in the S-HiAI (high adaptive immunity) and S-LoAI/HiII (low adaptive/high innate immunity) subtypes while down-regulated in the S-LoAI (low adaptive immunity) subtype. *ZBED6* is a transcription factor repressing IGF-2 which imprints the innate immune memory of macrophages [48]. *ZBED6* was down-regulated in the S-HiAI subtype, up-regulated in the S-LoII (low innate immunity) and S-LoAI (low adaptive immunity) subtypes.

By comparing up- and down-regulated DEG sets of each subtype, we observe that there is a significant overlap between the down-regulated genes in the S-HiAI subtype and up-regulated genes in the S-LoII (Pvalue= 5.2e-182) and S-LoAI subtypes (Pvalue= 5.4e-31). Similarly, down-regulated genes in the S-HiAI/LoII subtype had significant overlap with down-regulated genes in the S-LoII (Pvalue=5.5e-29) and S-LoAI/HiII subtypes (Pvalue=1.6e-24). (Figure 8). This highlights that some DEGs show opposite regulation across subtypes. This opposite patterning of DEGs and enriched pathways across subtypes may indicate response heterogeneity to the same intervention, some activating certain pathways while others suppress the same pathways.

**Figure 8.**
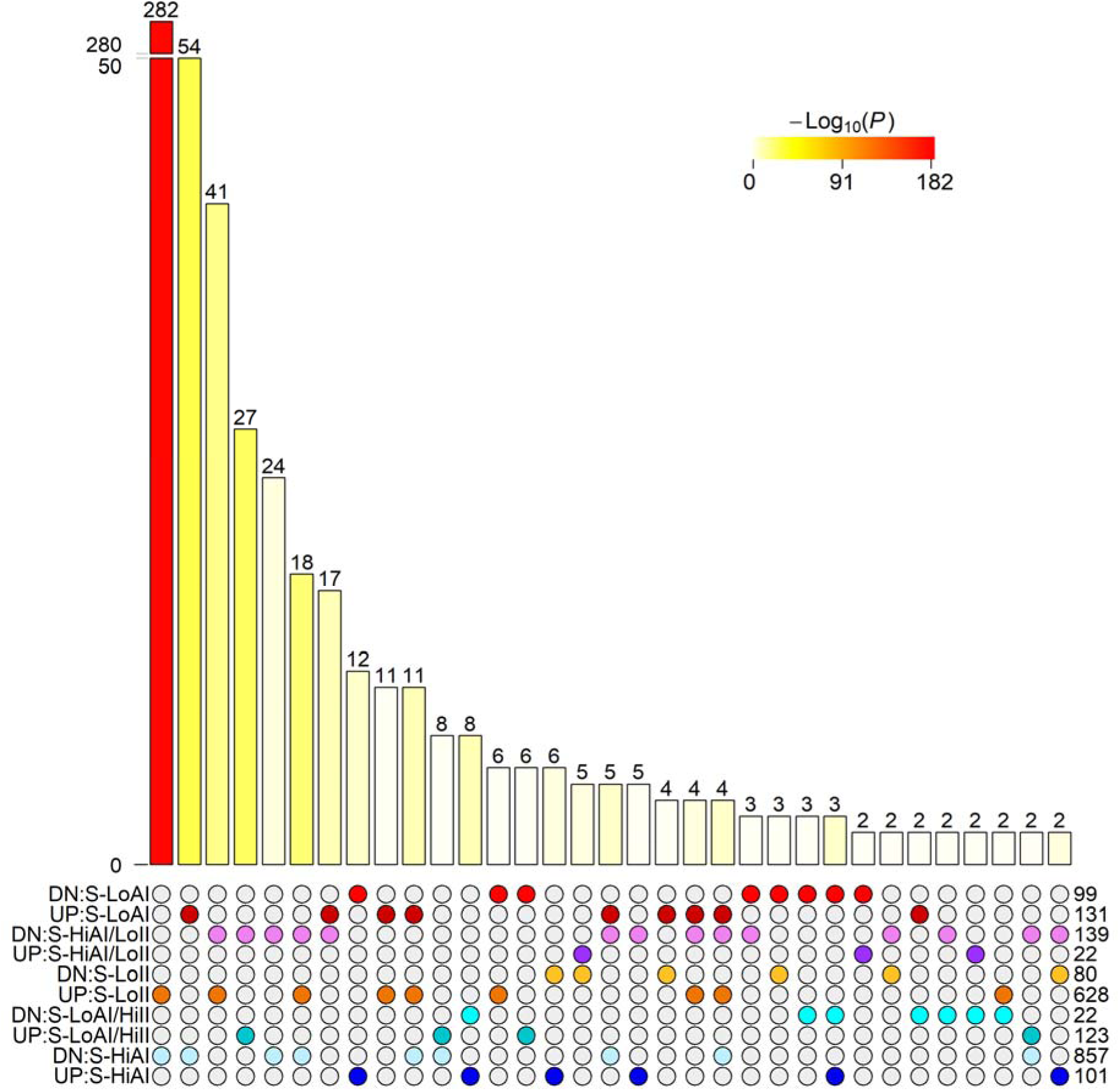
Intersection of differentially expressed gene sets by direction (up- and down-regulated). Overlapping number of genes across all combinations of five baseline molecular subtypes are shown. A colored circle indicates presence of the gene sets in each intersection. Number of genes in each gene set is shown on the right. In the gene set names, DN indicates the down-regulated DEGs of the indicated BMS compared to average of other four BMSs, and UP indicates the up-regulated DEGs (e.g., “UP: S-LoII” represents up-regulated genes in the low innate immunity subtype and “DN:S-HiAI” represents down-regulated genes in the high adaptive immunity subtype). The bars represent the overlap significance with color intensity representing the significance. The numbers shown on top of the bars indicate the overlapping number of DEGs.

### 3.4 Top molecular features of the predicted subtypes

Although the DEGs of each subtype showed significant associations with the subtypes, to have a complete understanding of subtype-specific molecular features, we identified the top subtype-specific molecular features by using the Kruskal-Wallis one-way ANOVA test. By comparing Z-transformed expression levels in each subtype to the other four subtypes (i.e., one vs others), we determined the top 10 features associated with each subtype. In the S-HiAI and S-LoII subtypes, the top 10 features were from transcriptomics and metabolomics data, in the S-LoAI/HiII and S-LoAI subtypes, they were all from metabolomics data, and, lastly, in the S-HiAI/LoII subtype, they were from glycomics and proteomics data (Figure 9A). Kynurenic acid, a metabolite linked to immune system regulation, was one of the top features of the S-LoAI/HiII (low adaptive/high innate immunity) subtype. Previously, a decrease in tryptophan along with an increase in kynurenine was found to be associated with regulation of inflammation and immunity in COVID-19 patients [49, 50]. Another immune response related metabolite, Creatinine, was one of the top molecular features of the S-LoII (low innate immunity) subtype [51]. C4b-binding protein alpha chain (*C4BPA*) protein controls the complement activation and was significantly associated with the S-HiAI/LoII (high adaptive/low innate immunity) subtype.

**Figure 9.**
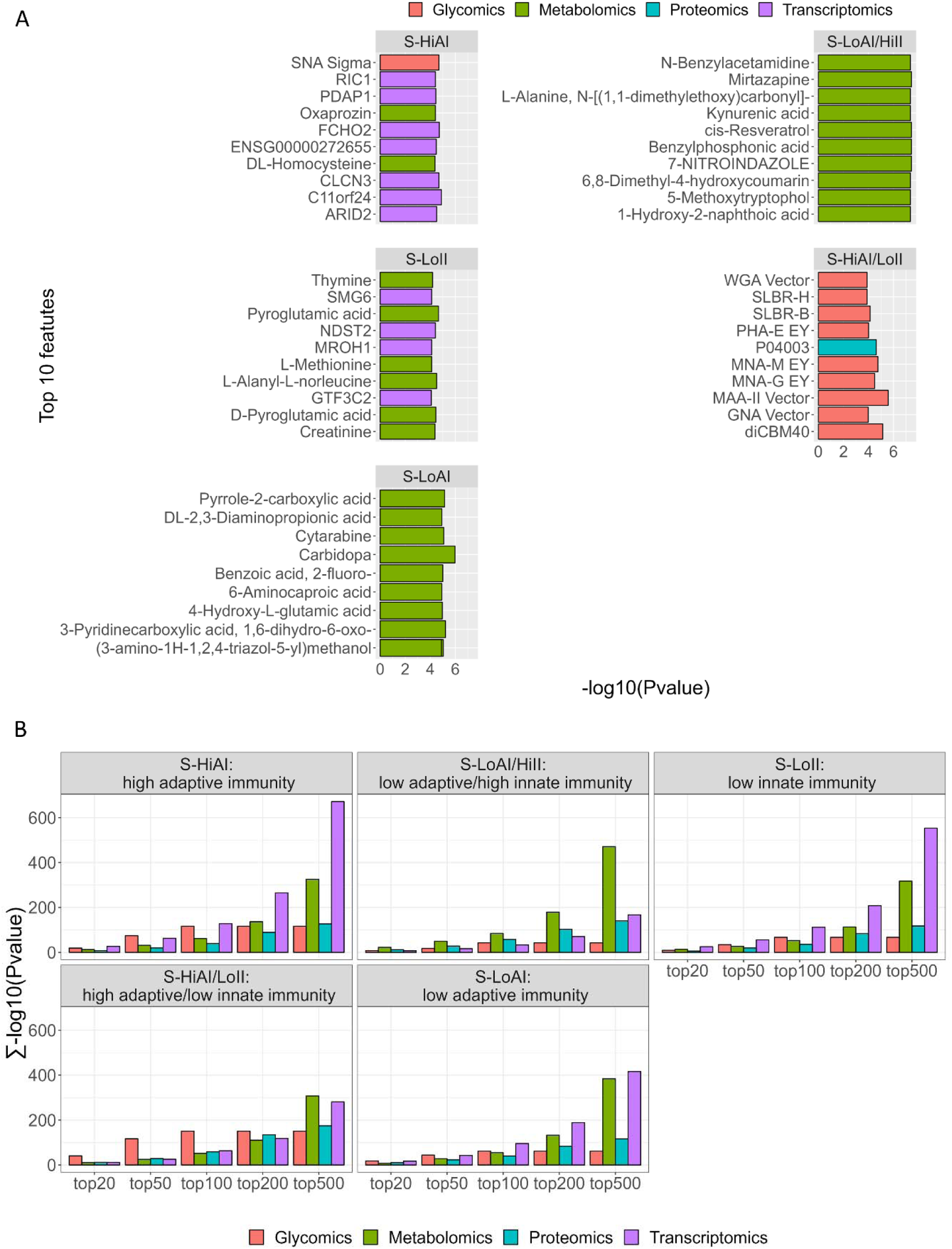
Top molecular features of the predicted baseline molecular subtypes. (A) Top 10 most significant molecular features of each BMS compared to the other BMSs (one vs others). Each bar represents one of the top 10 molecular features. (B) Cumulative significance of top 20, 50, 100, 200, and 500 features.

To demonstrate the contributions of each omic modality to each subtype, we calculated the cumulative significance of molecular features (Figure 9B). Notably, among the top 500 features, transcriptomics data had the most significant contribution in the S-HiAI and S-LoII subtypes, metabolomics in the S-LoAI/HiII, and both transcriptomics and metabolomics in the S-LoAI and S-HiAI/LoII subtypes. It is also important to note that, the number of molecular features were significantly different in different data modalities: 12,583 genes in the transcriptomics data, 15,903 metabolites in the metabolomics data, 273 proteins in the proteomics data, and 93 glycan binding lectins in the glycomics data. When we consider the top 100 features, glycomics data was the most significant contributing data to the S-HiAI/LoII subtype.

We also generated volcano plots of metabolite and protein expression values, as well as lectin microarray data, showing top differentially molecules of each subtype compared to the other four subtypes. N1-acetylspermidine was significantly up-regulated in the S-HiAI (high adaptive immunity) subtype. Up-regulated N1-acetylspermidine was previously associated with favorable outcome in severe COVID-19 patients and shown to have anti-inflammatory and immunostimulatory effects [50]. Dehydroepiandrosterone (DHEA, 5-Androsten-3beta-17-one) has been known to regulate immune function and was among the top down-regulated metabolites in the S-HiAI subtype [52]. Mangiferin, a natural flavonoid compound with anti-inflammatory and anti-oxidant properties, and Evodiamine, a supplement with anti-tumor and anti-inflammatory pharmacological effects, were both up-regulated in the S-LoAI/HiII (low adaptive/high innate immunity) subtype. Pyroglutamic acid, a cancer biomarker and previously shown as being elevated in the serum of ulcerative colitis patients, was found up-regulated in the S-LoII (low innate immunity) subtype [53]. Further, the up-regulated metabolite Pyroglutamic acid was found to be significantly altered in influenza high-risk groups in a recent study [10].

At the protein level, immunoglobulins were found to be down-regulated in the S-HiAI subtype while they were up-regulated at the mRNA level. This uncorrelated behavior could be due to various layers of regulation. The proteomics data is from serum while the transcriptomics data is from blood cells. Immunoglobulins could be kept on blood cells rather than being secreted into the serum. Alternatively, there could be a delay between transcription and translation or there could be post-transcriptional regulation that is reducing translation of the mRNAs into proteins. A similar pattern was observed in the S-LoAI/HiII subtype but in the opposite direction where immunoglobulins were down-regulated at the mRNA level while they were up-regulated at the protein level. Again, this could be due to several reasons. For example, there can be a delay between mRNA transcription and protein translation/secretion. Hence, recent transcriptional activity may be decreased (showing lowered mRNA levels) while protein levels reflect accumulation of immunoglobulins produced over a longer period. Immunoglobulins in the serum may be coming from other tissues, and the blood transcriptomics data may not be capturing these.

Despite the smaller probe set, we observed subtype-specific differentially expressed lectin microarrays. Notably, anti D-dimer, anti sialyl Lewis X, and anti Lewis X were up-regulated in the S-HiAI (high adaptive immunity with up-regulated immunoglobulin production and B cells) subtype, while being down-regulated in the S-LoII (low innate immunity with down-regulated immunoglobulin production, and complement activation) subtype. In a recent study, baseline Lewis A antigen (Le^a^) levels were found to be significantly higher in influenza vaccine non-responders compared to responders [8]. Follow-up glycoproteomic analysis showed that Le^a^-bearing proteins were enriched in complement activation pathways indicating a potential role for glycosylation in mediating complement.

### 3.5 Clinical and demographic features of the predicted subtypes

To investigate the clinical characteristics of the predicted subtypes, we examined the relationship between demographic and clinical features of the cohort and subtype groups. The subtypes were not associated with sex, age, BMI, pre-vaccination status, or month of vaccination based on the Kruskal-Wallis one-way ANOVA test.

## 4 Discussion

Immunological responses to vaccination are highly heterogeneous between individuals. To explore the molecular underpinnings of this variability, we performed an integrative multi-omics subtyping analysis in a cohort vaccinated against influenza.

In contrast to previous vaccination studies that focused on associations between vaccination response and physiological factors, our approach profiles molecular differences between individuals independently of physiological and clinical variation. We incorporated four different baseline omic datasets collected from 62 individuals of Caucasian ancestry. While revealing potential molecular signatures of vaccination response, the limited diversity warrants caution in generalizing our findings at this stage. Extending this integrative analysis to a larger and more inclusive cohort is needed in future studies.

Our study highlights the potential value of multi-omic subtyping in exploring molecular mechanisms underlying immune response heterogeneity. We observed opposite regulation of adaptive and innate immune pathways in different subtypes. For example, adaptive immunity was up-regulated in the S-HiAI subtype yet down-regulated in the S-LoAI subtype. Additionally, up-regulated innate immune response genes accompanied by down-regulated adaptive immune response genes characterized the S-LoAI/HiII subtype, while the opposite pattern characterized the S-HiAI/LoII subtype. Collectively, these findings indicate subtype-linked immune orientation.

Investigation of potential associations between predicted subtypes and pre- and post-vaccination antibody levels yielded significant differences across subtypes despite the absence of significant changes in antibody levels post-vaccination. For example, the average IgA and A/H3N2 IgA levels were higher at baseline in the S-LoAI/HiII subtype (pre-existing low adaptive/high innate immunity) and lower in the S-HiAI subtype (pre-existing high adaptive immunity), relative to other subtypes. This subtype-linked variation in IgA persisted post-vaccination. These results suggest that heterogeneity in vaccine responses is strongly impacted by baseline immune status.

Different omics modalities had variable contributions across the subtypes. Transcriptomic and metabolomic data predominated in four of the five subtypes, potentially reflecting the greater depth of these datasets—RNA sequencing assayed 12,583 genes and metabolomics quantified 15,903 metabolites, compared to just 273 proteins and 93 glycan binding lectins for the proteomic and glycomic data respectively. Notably, when considering only the 100 most discriminative features, glycomic data were most prevalent in defining the S-HiAI/LoII subtype. Thus, the predictive power of the individual omics modalities should not be underestimated, despite differences in scale. Overall, our integrated multi-omics approach leverages complementary contributions of the diverse dataset components, providing subtype predictions enriched in different biological pathways.

We observed heterogeneity in the immunoglobulin response across subtypes, i.e. in one subtype, immunoglobulins were upregulated at the protein level but downregulated at the mRNA level, while the opposite pattern was seen in another subtype. It indicates a disconnect between mRNA and protein regulation for immunoglobulins within the subtypes. This could be driven by factors such as differences in transcription/translation timing, source cell, post-transcriptional regulatory mechanisms, and kinetic rates between mRNA production and protein secretion. The opposite regulation of immunoglobulins at mRNA and protein level between subtypes on the other hand, highlights heterogeneity in the immunoglobulin response across subtypes.

Previous studies report that pre-existing immunity might play a role on vaccine effectiveness [54]. Repeated influenza vaccination may enhance or attenuate effectiveness [55]. Another potential impact on immunological memory is previous infections [56]. Our pathway enrichment analysis revealed two subtype groups (S-HiAI and S-HiAI/LoII) distinguished by up-regulation of the adaptive immune response pathway. More specifically, B-cell mediated immunity pathways were enriched in the S-HiAI subtype, whereas T-cell mediated immunity predominated in the S-HiAI/LoII subtype. Additionally, we observed subtype-specific regulation of innate immune response, with up-regulation of antiviral defense pathways occurring exclusively in the S-LoAI/HiII subtype and down-regulation in the S-HiAI/LoII subtype. Our findings corroborate previous studies indicating immunological memory is shaped by vaccination history and prior infections, and that it has a major role in the response to vaccination. Future research exploring how repeated vaccination and past infection alter gene expression profiles pre- and post-vaccination would extend our understanding of the mechanisms underlying diverse vaccine responses.

## Funding

This work was supported in part by the Center for Influenza Vaccine Research for High Risk Populations (CIVR-HRP) HHS-NIH-NIAID-BAA2018, by CRIPT (Center for Research on Influenza Pathogenesuis and transmission), a NIAID-funded Center of Excellence for Influenza Research and Response (CEIRR, contract # 75N93021C00014) and by NIAID grant U19AI168631 to A-GS, and by the Division of Intramural Research at NIAID/NIH.

## Conflict of Interest

The A.G.-S. laboratory has received research support from GSK, Pfizer, Senhwa Biosciences, Kenall Manufacturing, Blade Therapeutics, Avimex, Johnson & Johnson, Dynavax, 7Hills Pharma, Pharmamar, ImmunityBio, Accurius, Nanocomposix, Hexamer, N-fold LLC, Model Medicines, Atea Pharma, Applied Biological Laboratories and Merck, outside of the reported work. A.G.-S. has consulting agreements for the following companies involving cash and/or stock: Castlevax, Amovir, Vivaldi Biosciences, Contrafect, 7Hills Pharma, Avimex, Pagoda, Accurius, Esperovax, Farmak, Applied Biological Laboratories, Pharmamar, CureLab Oncology, CureLab Veterinary, Synairgen, Paratus, Pfizer and Prosetta, outside of the reported work. A.G.-S. has been an invited speaker in meeting events organized by Seqirus, Janssen, Abbott and Astrazeneca. A.G.-S. is inventor on patents and patent applications on the use of antivirals and vaccines for the treatment and prevention of virus infections and cancer, owned by the Icahn School of Medicine at Mount Sinai, New York, outside of the reported work. All other authors declare no conflict of interest.

## Author Contributions

CSB, CF, and BZ contributed to the conception and design of this study. DJ, DG, CV, LM, EG, TR generated the data. CSB performed the analysis and wrote the manuscript with input from all authors. All authors read and approved the manuscript.

## Data Availability Statement

We deposited the combined multi-omic data from 62 influenza vaccine recipients to https://www.synapse.org/#!Synapse:syn52749029.

